# Nonrandom spatial distribution of Neotropic Cormorants (*Phalacrocorax brasilianus*) along a coastal highway in Lima, Peru

**DOI:** 10.1101/2020.11.11.377903

**Authors:** Sebastián Lozano-Sanllehi, Carlos B. Zavalaga

## Abstract

Neotropic Cormorants (*Phalacrocorax brasilianus*) are common seabirds along the Peruvian coast. They frequently perch on trees, poles and port structures in urban areas, causing discomfort and esthetic problems due to the dropping of their feces on infrastructure and people. Hundreds of these birds rest on lighting poles and telephone cables along a 12.7 km highway in the coastal strip of the city of Lima, Peru. We hypothesized that the distribution of the cormorants along this highway is clustered and could be associated with physical features of both the coast and the adjacent marine area. Half-monthly or monthly surveys were performed from July 2018 to March 2020 in the Circuito de Playas de la Costa Verde highway. At each survey, cormorants were counted per lighting pole and adjacent telephone cables (pole-cable) at four count hours (0600 h, 1000 h, 1400 h and 1800 h). Our results revealed that daily bird numbers varied from 46 to 457 individuals and that only 17% of the total number of pole-cables (N = 651) was occupied once by at least one individual. The number of cormorants also varied between count hours within the same day (higher numbers at 1000 h and 1400 h). Birds were clustered into a maximum of five hotspots along the highway. According to the Akaike’s information-theoretic approach applied to Poisson GLMM, higher numbers of cormorants on pole-cables were associated mainly to a closer distance from these structures to the shoreline and to the surf zone, suggesting that Neotropic Cormorants may select such pole-cables as optimal sites for sighting and receiving clues of prey availability. Based on the results, the use of nonlethal deterrents and the relocation of these birds to other perching structures on nearby groynes could be the most suitable and eco-friendly solution for the problems caused by their droppings.

## Introduction

Overfishing, bycatch, habitat destruction, introduction of invasive species, hunting and pollution, among others, have caused both drastic decreases in the number and changes in the distribution of various species of seabirds worldwide [1, 2], many of which (42%) are considered in some extinction category [3, 4]. Conversely, other species of seabirds have been favored by the increase in available food in fishing discards [5–7], the presence of coastal landfills [8, 9] and the aquaculture industry [10, 11]. This has caused seabirds to interact more frequently with humans and in some cases to be considered pests [12, 13]. It has been shown that some species of seagulls can spread diseases [14–16], transport pollutants [17], damage urban infrastructure [18, 19], and collide with airplanes [20, 21]. Some species of cormorants, on the other hand, obtain their food from fish farms [22, 23] and generate conflicts with fisheries [24]. In Peru, the droppings of cormorants, among other species of seabirds, cause deterioration in port facilities [25]. In coastal cities of Chile, the excretions of Neotropic Cormorants (*Phalacrocorax brasilianus*, hereinafter “NECO”) present in trees and public lighting poles cause nuisance to people and damage to infrastructure and vegetation of parks and avenues [26]. A similar problem with this species occurs in Costa Verde, the coastal strip of the Miraflores bay in the city of Lima, Peru [27], through which the Circuito de Playas de la Costa Verde highway (CPCV) is extended.

The NECO is a species of wide distribution in the American continent, from the southern United States to Cape Horn, Chile [28, 29]. It is present in a wide variety of coastal, Amazonian and high Andean ecosystems [30–32], from sea level to altitudes up to 4800 m.a.s.l. [28, 33]. It feeds in shallow waters [34, 35], both in continental and marine water bodies [29, 36], and its diet is composed of a large variety of prey, mainly benthic fish [37–39]. It frequents places with perching structures such as trees, poles, cables, buoys or rocks, where it performs activities of preening, feather drying and daytime rest [40, 41].

The first records of the presence of NECOs occupying tall trees and light poles at the top of the cliffs of Costa Verde date from the mid-1950s [42]. However, over the years, the construction of vehicular roads, beaches, rock groynes and other infrastructure has led these birds to use public lighting poles and telephone cables present along the CPCV to rest and preen, thus favoring the contact of their feces with the road, cars, infrastructure and passers-by [27, 43]. It is speculated that their droppings could cause public health problems due to their possible content of pathogens [44], as happens in other birds [45]. The droppings could also cause traffic accidents when falling on the windshield of cars, or corrosion to vehicles and infrastructure, such as occurs with the Double-crested Cormorant (*Phalacrocorax auritus*) in the Columbia River (Astoria) Bridge, United States [46, 47]. Just as it happened in the city of Antofagasta, Chile [26], these problems could intensify or move to other areas over the years because Costa Verde is subject to a growing implementation of urban projects.

Currently, the absence of solutions to this environmental problem brings with it the need to evaluate and establish eco-friendly proposals, which must be included in the future design of projects in Costa Verde. There are mitigation measures used for NECOs in Arica and Iquique, Chile, which have had variable results; these include the relocation of colonies, the destruction of nests, the elimination of individuals, the use of sound and visual stressors, among others [26]. In the province of Chubut, Argentina, the elimination of NECOs and the harassment of their colonies are used as methods against presumed impacts of this species in fishing and aquaculture activities [48]. For the Peruvian case, to determine which management proposals are the most appropriate, it is important first to know how and why these birds are present along the CPCV.

In this study, half-monthly and monthly counts were performed between July 2018 and March 2020 on poles and telephone cables that these birds used as perches along the CPCV. The objectives were (1) to evaluate the spatial distribution and temporal variation in the number of NECOs in the CPCV, and (2) to examine the influence of physical features of both Costa Verde and the adjacent marine area in their spatial distribution. Based on the results, eco-friendly solutions to the environmental problem caused by their excreta were proposed. This is the first study aimed at solving problems caused by seabirds in urban-coastal areas of Peru.

## Methods

### Study area

This study was carried out in Costa Verde, the coastal strip of the Miraflores bay in the city of Lima, Peru (Fig 1). This is a tourist-recreational space with a large influx of vehicles and people throughout the year, especially in summer. It has beaches, parks, sports complexes, restaurants, clubs, parking areas adjacent to the beaches and infrastructure for pedestrian and vehicular traffic (Fig 2A and 2B). Along Costa Verde, the CPCV highway extends, which is limited to the east by cliffs up to 100 m high and to the west by sandy, gravel and pebble beaches, usually narrow, with rock groynes perpendicular to the coastline ([43], Fig 2A). The section of this road that was assessed in the study includes from the entrance to Club de Regatas “Lima” (club) from the south (12°10’0.2”S, 77°1’48”W) to Bajada Escardó (access road) from the north (12°5’14.4”S, 77°5’38.8”W; Fig 1). This section is 12.7 km long and has two to three lanes for vehicular traffic on both the outward and return tracks (Fig 2C). It also presents public lighting poles and telephone cables that favor the perch of NECOs and other birds such as gulls and black vultures (*Coragyps atratus*, Fig 2D).

**Fig 1.**
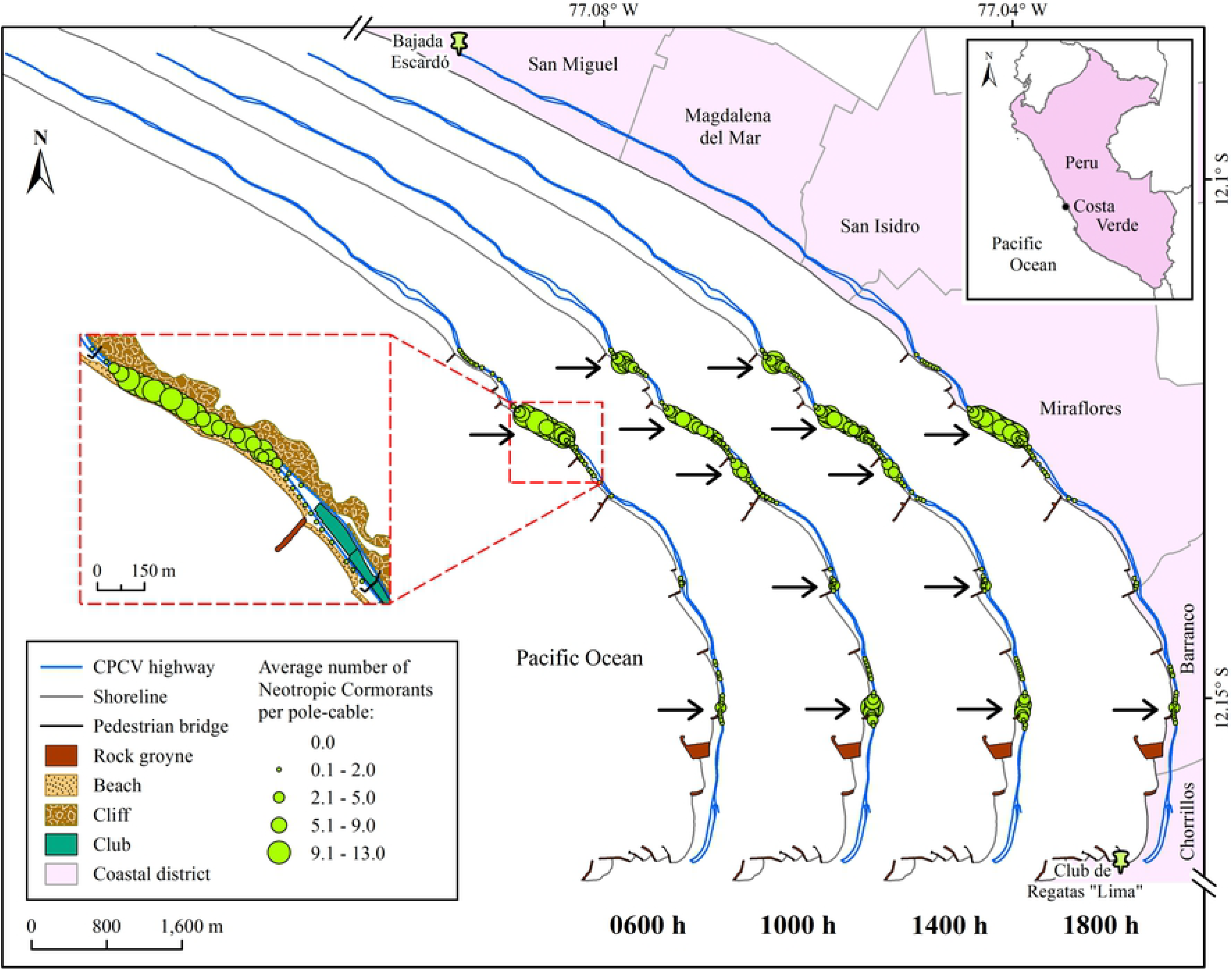
Location of the study area and spatial distribution of the Neotropic Cormorants (*Phalacrocorax brasilianus*) by count hour in the Circuito de Playas de la Costa Verde highway in Lima, Peru. The size of the green circles is in function to the study-period average number of cormorants per pole-cable. The arrows indicate the aggregation hotspots of the birds.

**Fig 2.**
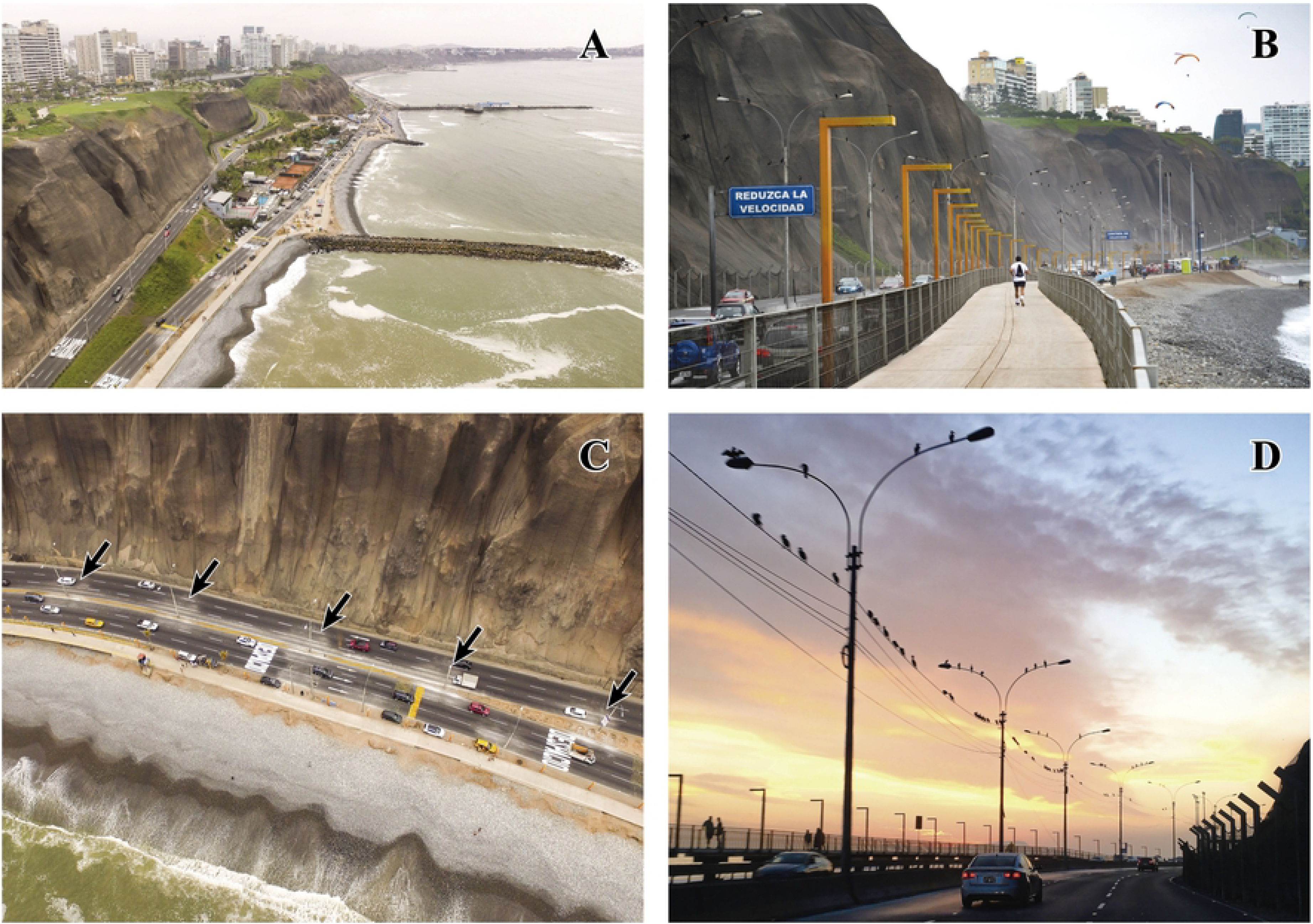
Features of Costa Verde. (A) Cliffs, clubs, groynes and pebble beaches. (B) Infrastructure for pedestrians. (C) Vehicular road with presence of cormorant feces around each lighting pole (pointed by black arrows). (D) Neotropic Cormorants (*Phalacrocorax brasilianus*) perching on lighting poles and telephone cables.

Using ArcGIS 10.5 software [49] and based on World Imagery images (projected into UTM zone 18S) from the ArcGIS Online service, different attributes of Costa Verde were digitized, such as public lighting poles, vehicular roads, pedestrian bridges, cliffs, groynes, beaches and the shoreline. This allowed the calculation of the position and distance of different attributes required in the data analysis. Each lighting pole was identified with a unique code.

### Counts

Visits to the CPCV were made half-monthly (in the middle and at the end of each month), from July 14, 2018 to June 30, 2019, and monthly (at the end of each month), between July 31, 2019 and March 27, 2020. In both cases, four surveys were performed in the day of count: 0600 h, 1000 h, 1400 h and 1800 h, resulting in a total of 96 half-monthly and 33 monthly counts (on March 27, 2020, NECOs were only counted at 1000 h). The 0600 h counts were performed at dawn (0530-0630 h), as soon as the daylight was sufficient to count the NECOs and thus record their number and distribution before they began to depart. In the case of 1800 h, the counts were performed as late as possible (1740-1840 h) before darkness; this was done to record their number and distribution upon returning to their place of overnight roosting.

The number of NECOs in the CPCV was determined by direct count of birds recorded in high-resolution videos. For this, a GoPro HERO6 camera (4K/60fps) was attached to the roof of an automobile that moved at a speed <30 km/h. The camera lens pointed to the top of the poles and cables. Beaches and other surrounding areas were not included in the counts since they were not occupied by the NECOs during the study period. The counting of individuals was performed per lighting pole and that number was associated with the code of the pole. In the case of the NECOs present in telephone cables, the length of the cable between poles was divided into two equal parts, and the NECOs on each half were assigned to the nearest pole. Therefore, the term “pole-cable” is used hereinafter as a unit of counting and analysis. The Franklin’s Gull (*Leucophaeus pipixcan*), a seasonal migratory and very abundant species, as well as other species of birds present in fewer numbers on pole-cables, were also counted.

There were no specific permits and approvals required for data collection at field since Costa Verde is a public area and no animal experimentation was performed.

### Definition and measurement of explanatory variables

The distribution of poles along 12.7 km of the CPCV was regular. Neither the poles nor the telephone cables had deterrents that prevented birds from occupying them, such as spikes or bird wires. The number of NECOs on pole-cables was examined by variables of physical features of both Costa Verde and the adjacent marine area (<1 km from the shoreline), which could be related to the foraging behavior of this seabird species and the accessibility to their prey.

#### Distance from the pole-cables to the shoreline (DS)

The distance from the pole-cables to the shoreline varies along the CPCV. Prior to this study, it was observed on several occasions that in sections of the highway closer to the shore there was a greater number of NECOs on pole-cables. Thus, to examine this effect, the minimum distance from each pole-cable to the shoreline (line that passes between the high and low tide marks) was measured in ArcGIS.

#### Distance from the pole-cables to the surf zone (DSZ)

The surf zone is the relatively narrow strip that borders the ocean beaches and covers from the shoreline to the last breaking wave offshore [50]. It is a very dynamic and productive zone, as it holds a wide variety of life forms [51]. In Costa Verde, its amplitude varies along the coast and could represent areas preferred by the NECOs for their foraging activities.

Five satellite images of Costa Verde obtained from the historical images of Google Earth Pro 7.3 software [52] were used with dates coinciding with the study period (October and December 2018, March and April 2019, February 2020), which were georeferenced in ArcGIS. On each image, a layer of the shoreline sections in front of a surf zone with an amplitude greater than 70 m (measured from the shoreline to the last breaking wave close to the coast) was digitized. Lower amplitudes were not taken into account because they were common along the coastline. Once the layers of the five images were generated, they were overlapped. The sections with matches in at least three images were defined as the surf zone sections. Finally, the minimum distance from each pole-cable to the nearest surf zone section was measured.

#### Distance from the pole-cables to the nearest groyne (DG)

Coastal protection structures alter the hydrodynamic regimes and depositional processes of the coastal zone [53], which can lead to changes in the habitat, composition and distribution of marine species [54]. Thus, the presence of rock groynes in Costa Verde could generate feeding areas preferred by the NECOs. For the calculation of this variable, the minimum distance from each pole-cable to the base of the nearest groyne was measured in ArcGIS.

#### Perimeter of the nearest groyne (PG)

The hard substrate of groynes allows many marine organisms to settle [54, 55]. Thus, the larger groynes and whose contour has more contact with the seawater would generate a greater feeding area for the NECOs. For this reason, each pole-cable was assigned the length of the contour (in contact with the seawater) of the nearest groyne.

#### Distance from the shoreline to the 7 m isobath (D7)

According to various authors, NECOs have a preference for areas near the coast [28], where they feed in shallow waters close to the shoreline [34, 48, 56]. In Costa Verde, their number on pole-cables could be associated with a preference for shallow marine areas, which are more extensive at a greater distance from the shoreline to the isobaths.

The isobaths of the marine area of Costa Verde were obtained from the digitized nautical chart No. 2237, corresponding to the Miraflores bay [57]. The minimum distance from the shoreline (in front of each pole-cable) to the 7 m isobath was measured in ArcGIS. The choice of this isobath was based on records of the preference of NECOs to feed in waters with depths less than 10 m [35, 58, 59].

#### Transparency of seawater (T)

In general, seabirds tend to congregate in areas of high prey density [60]. However, its accessibility to these depends on variables such as water clarity [61]. Therefore, possible changes in the transparency of seawater could influence the presence of NECOs on pole-cables in the CPCV.

This variable was measured between the months of October 2018 and January 2019 (one measurement per month) on dates coinciding with surveys. These measurements were performed with a Secchi disk from a boat in areas with depths between 5 and 8 m and at an approximate minimum distance of 500 m from the shoreline. The presence of waves prevented going further inshore. Data were taken at the beginning, center and end of transects of 400 m perpendicular to the shoreline, in front of five sections with presence and five sections with absence of NECOs (each section of 15 pole-cables). Finally, the values obtained by transect were averaged, thus obtaining a total of 40 data points of seawater transparency throughout the evaluation period of this variable.

### Analysis of the spatial distribution and temporal variation of the number of NECOs

Count data were examined to determine NECOs’ random or cluster distribution along the CPCV and to identify occurrence hotspots. For management purposes, the number of NECOs was analyzed by district (Callao, San Miguel, Magdalena del Mar, San Isidro, Miraflores, Barranco and Chorrillos). The district boundaries layer was obtained from the GEO GPS PERÚ website [62].

For each count hour (0600 h, 1000 h, 1400 h and 1800 h), the number of NECOs per pole-cable of all counts was averaged. These averages were processed in ArcGIS to generate maps and define the aggregation “hotspots” of the NECOs. A set of consecutive pole-cables with two or more NECOs on each one was considered as a “hotspot”.

To determine whether the spatial distribution of the NECOs was random or clustered, the Moran spatial autocorrelation measure (global Moran’s I) was applied to the averaged data of each count hour using ArcGIS. Likewise, frequency graphs of the number of NECOs per pole-cable and the two-sample Kolmogorov-Smirnov (KS) test (*stats* R package [63]) were used to assess and compare the level of aggregation between count hours.

The temporal variation of the number of NECOs among count hours (0600 h, 1000 h, 14000 h and 1800 h) and among seasons (austral summer, autumn, winter and spring), and their interaction, were analyzed with a generalized linear model (GLM, *stats* R package [64]). Additionally, the correlation between the number of NECOs and the number of Franklin’s Gulls (daily maximum number of gulls and NECOs in the CPCV) was examined using another GLM (*rstatix* R package), including the year of count as a covariate.

Statistical analyses were performed in R 3.5.1 software [65] with a significance level of α = 0.05. The averages are expressed ± 1 SD.

### Correlations with explanatory variables

Collinearity between five of the six explanatory variables (DS, DSZ, DG, PG and D7) was examined using pairplots (two-way associations) and variance inflation factors (VIF, high-dimensional collinearity, *car* R package [66]). Covariation was assessed using the threshold of 0.65 for correlation coefficient and 3 for VIF. In order to reduce covariation between DG and DS (Pearson correlation, r = 0.8), and between DSZ and DS (r = 0.8), and based on the fact that both DG and DSZ included the most extra information in relation to DS, new variables were created controlling for DS (DG:DS ratio, DSZ:DS ratio). Because the two created variables were highly correlated (r = 0.92), none of them were used in the same model. D7 also showed high correlation in respect to the other variables excepting PG (r = 0.04), so it was analyzed separately. Finally, the variables from each set [(DS, DG:DS ratio, PG), (DS, DSZ:DS ratio, PG) and (D7, PG)] showed VIFs lower than 2.

A generalized linear mixed model (GLMM) analysis (*glmmTMB* R package [67]) with a Poisson distribution was conducted for the data of the number of NECOs per pole-cable at 1000 h as the response variable; this count hour was chosen because it registered higher numbers of NECOs during the day and because it was one of the count hours of NECOs’ diurnal activity with a higher occupancy of pole-cables. Different models were generated that included the explanatory variables, according to the approach of *ad hoc* hypotheses. The code of the pole was included as a random effect to cope with pseudoreplication since each pole-cable was measured repeatedly during the study period. To account for autocorrelation found in the residuals, a first-order autoregressive (AR1) covariance structure was included, with ‘code of the pole + 0’ as the design matrix and the date of count as the grouping factor. For each model, non-significant variables (p > 0.05) were excluded and the non-overdispersion assumption was checked. The choice of the most appropriate model was performed through an information-theoretic approach (Akaike information criterion - AIC), taking into account the differences between AIC values (Δ_*i*_) and the Akaike weights (ω_*i*_, [68]).

Unlike the five variables abovementioned, transparency of seawater (T) data were measured by section of the CPCV and not by pole-cable. Therefore, the correlation analysis was performed separately, using a logistic regression (logit link function, *stats* R package [69]) with the presence or absence of NECOs on pole-cables as the response variable. For that purpose, on the same day of data collection, the set of 20 pole-cables located perpendicular to each transect was categorized with a value of presence (1) or absence (0) of NECOs.

Correlations and other statistical analyses were performed in R software with a significance level of α = 0.05. The averages are expressed ± 1 SD.

## Results

In the section of the CPCV studied, 651 public lighting poles were identified, of which 47% were concentrated in only two districts (Miraflores and Barranco, S1 Table), since more than half of the total length of the CPCV corresponds to these districts (6.7 km). Of the total poles, the largest number were single arm (55%) and double arm (40%) poles, and were distributed throughout the six districts. In respect of the groynes, Miraflores held 7 of the 12 that exist throughout Costa Verde, followed by Barranco (three groynes) and Chorrillos (two groynes), while in the other districts, these rocky structures were absent (S1 Table, Fig 1). In addition to the NECOs and Franklin’s Gulls, other bird species were present on the pole-cables in smaller numbers during the study period. In all counts, a daily maximum of 69 Band-tailed Gulls (*Larus belcheri*), 13 Rock Doves (*Columba livia*), 11 Black Vultures (*Coragyps atratus*), five Kelp Gulls (*Larus dominicanus*) and three Zenaida Doves (*Zenaida* sp) were recorded in the CPCV. It must be taken into account that these quantities are minimal since these species could also be present on beaches or other habitats in Costa Verde.

### Spatial distribution and temporal variation of the number of NECOs

During the entire study period, the average number of NECOs of all the counts in the CPCV was 232.4 ± 106.7 individuals. The highest number of NECOs was recorded on November 27, 2019 (457 birds), and the lowest on February 28, 2019 (46 birds, Fig 3). Of the 651 pole-cables present on this highway, only 17% were occupied by at least one NECO, while the rest were always empty. These occupied pole-cables were located only in the districts of Miraflores (78%) and Barranco (22%).

**Fig 3.**
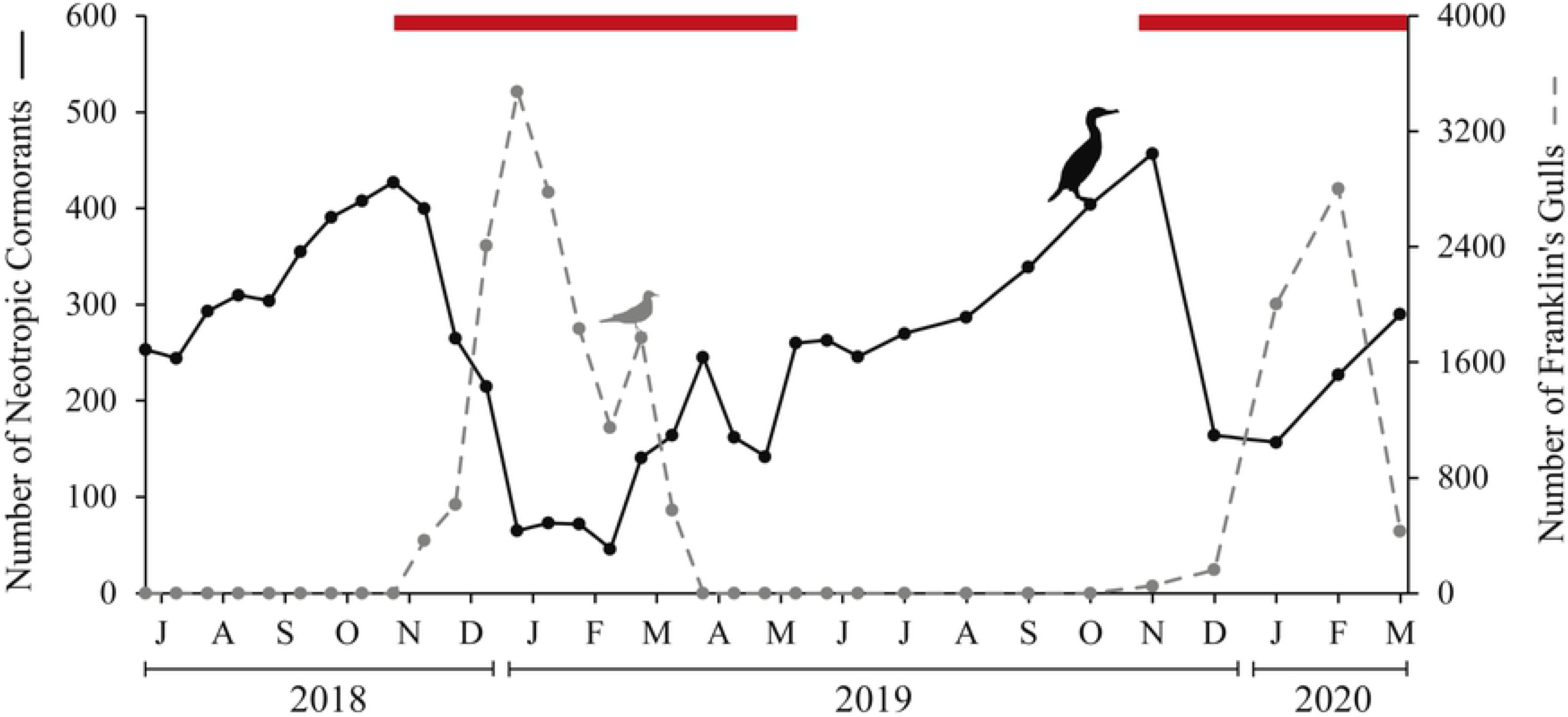
Temporal variation of the daily maximum number of Neotropic Cormorants (*Phalacrocorax brasilianus*) and Franklin’s Gulls (*Leucophaeus pipixcan*) in the Circuito de Playas de la Costa Verde highway in Lima, Peru. The red bars at the top of the graph corresponds to the breeding season of the Neotropic Cormorants, which was defined based on records in two coastal wetlands of the central zone of Peru (Los Pantanos de Villa [70] and El Paraíso [71]) and other studies that describe the duration of the stages of their reproductive cycle in different locations of the Peruvian coast [72–74].

#### Variations in spatial distribution

The spatial distribution of the NECOs was significantly clustered along the CPCV for each of the count hours: 0600 h (global Moran index, GMI = 0.439, z = 11.86, p < 0.001); 1000 h (GMI = 0.416, z = 11.13, p < 0.001); 1400 h (GMI = 0.41, z = 10.97, p < 0.001) and 1800 h (GMI = 0.438, z = 11.83, p < 0.001; Fig 1). At 0600 h and 1800 h, two hotspots were observed, located in Miraflores and Barranco, which gathered 93% of the total number of NECOs (Fig 1). In contrast, at 1000 h and 1400 h, the number of hotspots increased to five (four in Miraflores and one in Barranco) and gathered 84% of the total number of NECOs. The remaining percentage for the four count hours was distributed in other sections of the CPCV, in pole-cables with less than two NECOs on average (Fig 1). The differences in the number of hotspots reflect changes in the aggregation of NECOs per pole-cable. This aggregation was significantly different when comparing the count hours with two hotspots (0600 and 1800 h) with respect to the count hours with five hotspots (1000 h and 1400 h; KS test for each of the four paired comparisons, D ≥ 0.036, p < 0.001; Fig 4). At 0600 h and 1800 h, there were higher numbers of NECOs per pole-cable; more than 98% of the NECOs recorded in all counts during the study period (cumulative numbers, N_0600 h_ = 6,373 birds, N_1800 h_ = 6,838 birds) occupied pole-cables up to 22 individuals each (Fig 4A and 4D). In contrast, at 1000 h and 1400 h, more than 96% of these birds (cumulative numbers, N_1000 h_ = 7,825 birds, N_1400 h_ = 7,728 birds) occupied pole-cables with at most 14 individuals each, so the level of aggregation was lower (Fig 4B and 4C).

**Fig 4.**
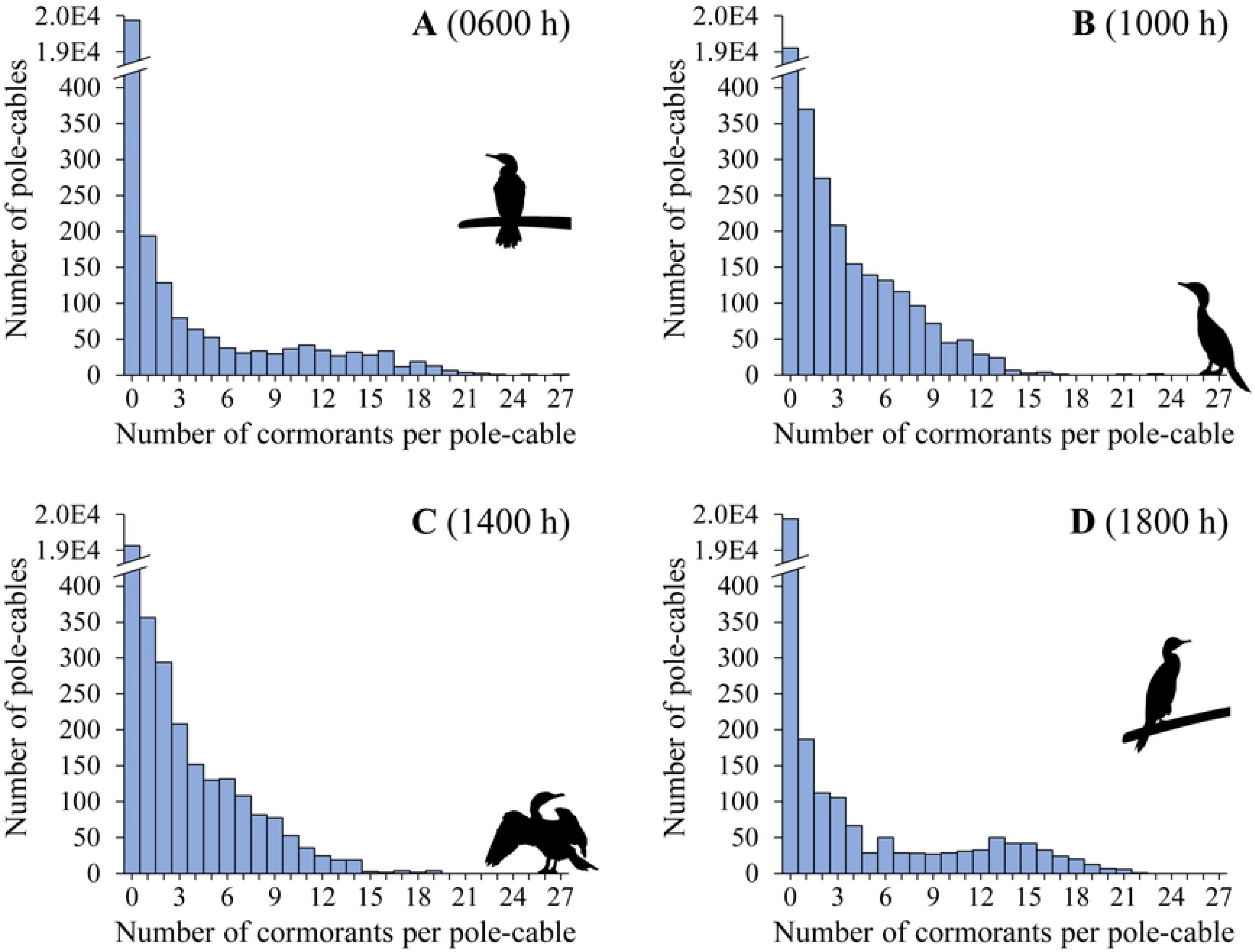
Frequency of the number of Neotropic Cormorants (*Phalacrocorax brasilianus*) per pole-cable, recorded from all surveys during the study period in the Circuito de Playas de la Costa Verde highway in Lima, Peru. (A) 0600 h. (B) 1000 h. (C) 1400 h. (D) 1800 h.

#### Temporal variation

The number of NECOs in the CPCV varied significantly between count hours within the same day (GLM, F_3_ = 3.46, p = 0.019), with the highest numbers of these birds at 1000 h (251.1 ± 110.7 individuals) and at 1400h (248.4 ± 111.6 individuals, Fig 5). On the other hand, the variation in the number of NECOs between seasons was also significant (GLM, F_3_ = 62.56, p < 0.001; Fig 5). In austral spring (Sep - Nov), a greater number of birds was observed in both 2018 and 2019. In summer (Jan - Feb) of 2019, the number of NECOs was the lowest recorded during the study period (Fig 3). The interaction between count hour and season was not significant (GLM, F_9_ = 0.15, p = 1), showing that the differences in the number of NECOs between count hours were maintained regardless of the season of the year (Fig 5).

**Fig 5.**
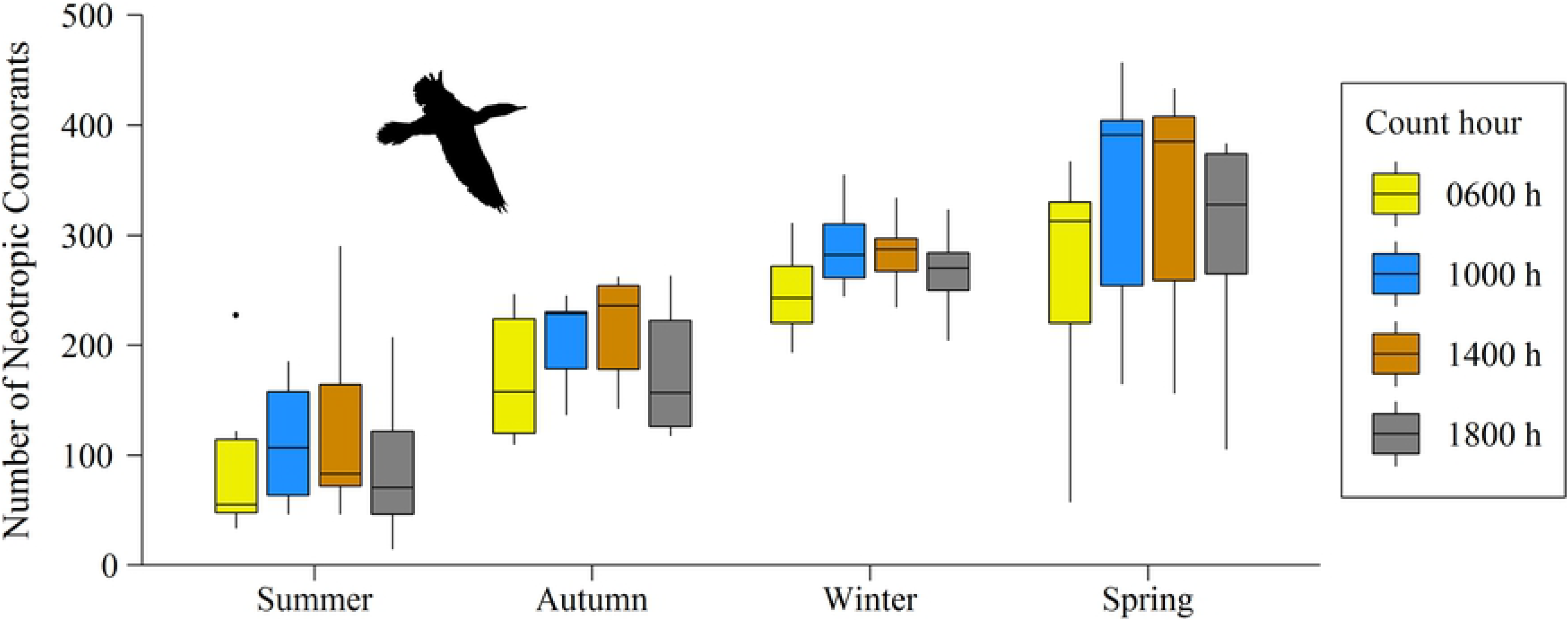
Temporal variation of the number of Neotropic Cormorants (*Phalacrocorax brasilianus*) by count hour and season of the year in the Circuito de Playas de la Costa Verde highway in Lima, Peru.

Franklin’s Gulls showed a typical pattern of a boreal migratory species. These birds were absent from the CPCV in the winter of 2018 and in the autumn and winter of 2019 (Fig 3). The first individuals began to appear in late November in both years, reaching peaks of 3,475 and 2,804 individuals during the summer (Jan - Mar) of 2019 and 2020, respectively. There was a significant inverse correlation between the number of NECOs and Franklin’s Gulls (GLM, F_1,30_ = 19.79, p < 0.001), with this relationship being similar between the period 2018-2019 and the period 2019-2020 (GLM, F_1,30_ = 1.98, p = 0.17, Fig 3). It should be noted that during the study period, the presence of Franklin’s Gulls was recorded on more than 90% of the pole-cables that were once occupied by a NECO (N = 111).

### Correlations with explanatory variables

The distances from the pole-cables to the shoreline (DS) ranged from 16 m to 290 m (Table 1), but the NECOs were located only in pole-cables at distances <60 m (Figure A in S1 File). The distances from the pole-cables to the surf zone (DSZ) presented a distribution very similar to the distances from the pole-cables to the nearest groyne (DG), with 50% of the data included between 18 and 400 m in both cases, and with NECOs perched on pole-cables at distances up to 280 m (DSZ, Figure B in S1 File) and 490 m (DG, Figure C in S1 File). On the other hand, 91% of the pole-cables were located near groynes with a perimeter between 87 and 270 m; the groynes of the remaining percentage (9%) had a perimeter greater than 635 m (Figure D in S1 File). The distance from the shoreline to the 7 m isobath (D7) had a multimodal behavior, with most of the values at approximately 445, 865 and 1215 m; in addition, almost 75% (N = 45) of the pole-cables occupied by NECOs were located in front of distances (D7) between 830 and 995 m, and the rest (N = 16 pole-cables) from 1195 m onwards (Figure E in S1 File). Finally, the transparency of seawater data (T) ranged from 74 cm to 515 cm depth (Table 1) and were distributed asymmetrically to the right, with 75% of the values between 74 and 208 cm depth (Figure F in S1 File); the T values associated with sections of the CPCV with presence of NECOs were also skewed to the right (Figure F in S1 File).

**Table 1.** Descriptive statistics of the variables used to explain the spatial distribution of the Neotropic Cormorants (*Phalacrocorax brasilianus*) in the Circuito de Playas de la Costa Verde highway in Lima, Peru.

| Variable | N | Mean ± SD | Range | Median |
| --- | --- | --- | --- | --- |
| DS (m) | 651 | 138.7 ± 79.9 | 15.8 - 289.6 | 132.3 |
| DSZ (m) | 651 | 1297.9 ± 1600.7 | 18.5 - 5615.8 | 396.5 |
| DG (m) | 651 | 1349.5 ± 1640.3 | 20.1 - 5719.3 | 386.7 |
| PG (m) | 12 | 259.8 ± 201.6 | 87.4 - 747.8 | 196.6 |
| D7 (m) | 651 | 796.6 ± 295.8 | 349.7 - 1254.6 | 843.9 |
| T (cm) | 40 | 200.4 ± 104.4 | 74.0 - 514.3 | 172.5 |

DS: distance from the pole-cables to the shoreline, DSZ: distance from the pole-cables to the surf zone, DG: distance from the pole-cables to the nearest groyne, PG: perimeter of the groyne, D7: distance from the shoreline to the 7 m isobath, T: transparency of seawater.

According to the Akaike’s information-theoretic approach, the models with individual variables had relatively high AIC values compared to the other models (Table 2), so they could not explain by themselves the spatial distribution of the NECOs in the CPCV. In contrast, the best model with lower AIC was M5 (DS [distance to shoreline] + DSZ:DS ratio [distance to surf zone:distance to shoreline ratio] + PG [perimeter of nearest groyne]; Table 2). The other proposed models presented AIC values differences greater than 2 with respect to the M5 model, so they were not taken into account. Additionally, the M5 model was given a higher Akaike weight (ω_*i*_ = 0.67) compared to the other models (Table 2), which reaffirms its choice.

**Table 2.** Poisson generalized linear mixed models to explain the spatial distribution of the Neotropic Cormorants (*Phalacrocorax brasilianus*) in the Circuito de Playas de la Costa Verde highway in Lima, Peru.

| Model | Description of the model | AIC | $\Delta_i$ | $\omega_i$ |
| --- | --- | --- | --- | --- |
| M5 | DS + DSZ:DS ratio + PG | 9861.3 | 0.0 | 0.67 |
| M4 | DS + DG:DS ratio | 9864.2 | 2.9 | 0.15 |
| M6 | DS + DSZ:DS ratio | 9864.3 | 3.0 | 0.15 |
| M1 | DS | 9867.7 | 6.4 | 0.03 |
| M2 | DSZ:DS ratio | 10046.8 | 185.5 | 0.0 |
| M3 | D7 | 10048.1 | 186.8 | 0.0 |

AIC: Akaike values, $\Delta_i$ : Akaike values differences, $\omega_i$ : Akaike weights, DS: distance from the pole-cables to the shoreline, DSZ: distance from the pole-cables to the surf zone, DG: distance from the pole-cables to the nearest groyne, PG: perimeter of the nearest groyne, D7: distance from the shoreline to the 7 m isobath.

According to the chosen model, the number of NECOs per pole-cable was significantly negatively related to DS (Poisson GLMM, β = −0.174 ± 0.017 SE, z = −10.38, p < 0.001), and to DSZ:DS ratio (β = −0.247 ± 0.099 SE, z = −2.497, p = 0.013; S2 Table). On the other hand, PG had also a significant but weak negative effect on the number of NECOs per pole-cable compared to the other variables (β = −0.003 ± 0.002 SE, z = −2.229, p = 0.026; S2 Table). It should be mentioned that, as a result of having created DSZ:DS ratio as a new variable to meet the assumption of non-collinearity, it remained highly correlated in respect to DSZ (r = 0.95, p < 0.001).

The correlation between the presence of NECOs on pole-cables and the transparency of seawater (T) was not significant (Logistic regression, *χ*^*2*^Wald = 0.11, p = 0.74).

## Discussion

Our results show that hundreds of NECOs congregate on pole-cables in a maximum of five hotspots along the CPCV highway, in the districts of Miraflores and Barranco.

### Presence of NECOs in the CPCV

In the coast of Peru, NECOs frequently occupy areas with artificial structures where they can perch and reproduce, such as metal beams under docks [75], metal ladders for access to breakwaters [25], wooden winches (structures to support pulleys) [31, 76], beams of load in ports [25], and public lighting poles [27]. They are also found in parks [77] and wetlands [32]. The presence of NECOs in Costa Verde is not a recent event, and their persistence trough years could be explained by the implementation of urban projects in Metropolitan Lima, including its coastal strip [78]. The construction of vehicular roads and touristic-recreational infrastructure has favored that these birds use structures such as pole-cables along the CPCV for resting [27]. Also, this permanency of the NECOs in Costa Verde suggests that the features of the Miraflores bay favor the presence of different prey from which these birds feed. It should be noted that possibly part of the population of NECOs has been displaced from nearby natural environments such as Pantanos de Villa, Humedales de Ventanilla (wetlands), Isla El Frontón and Isla San Lorenzo (islands), probably due to food shortage or loss of their habitat.

### Number of NECOs in the CPCV

The daily number of NECOs recorded in the CPCV during the study period ranged between 46 and 457 individuals. NECOs are also present in different guano islands and headlands along the coast of Peru, where the number of birds in 11 of these locations rarely exceeds 400 individuals (e.g., [75, 76, 79–81]). For example, between 1990 and 2016, the population of NECOs recorded in Islas Lobos de Afuera and Punta Salinas varied to a maximum of 450 and 472 birds, respectively [79, 82]. These numbers are very similar to those found in this study, which suggests that the CPCV colony is representative.

While the daily variations in the number of NECOs in the CPCV may respond to diurnal activity patterns related to their foraging behavior, its seasonal variations may be linked to their annual life cycle. The greatest numbers of NECOs on the pole-cables were found between mid-morning and mid-afternoon, suggesting that these birds spend most of the day on these structures resting or in plumage maintenance and drying activities [83–85]. Unlike other colonies [41, 83, 84], in the CPCV most of the NECOs kept roosting on the pole-cables during the nighttime. It is possible that a low percentage of these birds moved to other overnight places nearby the CPCV (e.g., boat structures in the Chorrillos dock), which could explain their lower number recorded at dawn and dusk. The marked variations in the number of NECOs between seasons in the CPCV seem to respond to the phenology of its annual life cycle. Low numbers of birds were recorded from the end of austral spring to the beginning of autumn, which corresponds to their breeding season in the central zone of the Peruvian coast [70, 71]. In addition, adult individuals with reproductive plumage were observed in the CPCV in November 2018 and December 2019, as well as several juveniles from November 2019 to March 2020 as a result of a possible good reproduction in 2019. The gradual increase of NECOs from the month of March onwards may be related to a progressive arrival of juveniles and adults from their breeding sites, as happens in other regions [41]. It is important to mention that one of these sites closest to Costa Verde is the wetland Pantanos de Villa [70], located 7-14 km away, so it is completely feasible that the birds of the CPCV move to that area and vice versa. The seasonal variations in the number of NECOs have also been observed in colonies of other regions, but with a different phenology [41, 84, 86, 87]. Another nonmutually excluding hypothesis that explains the seasonal variation in the number of NECOs in the CPCV is the strong inverse relationship with the number of Franklin’s Gulls. These boreal migratory seabirds arrive at the CPCV in mid-November, reaching a peak of 3,475-2,804 individuals between December and March, months in which the number of NECOs decreases considerably. Franklin’s Gulls make extensive use of pole-cables and occupy different sections of the CPCV, including those with presence of NECOs. Thus, there could be competition between both species for perching space on pole-cables.

### Variables related to the distribution patterns of the NECOs in the CPCV

This study has shown that higher numbers of NECOs on pole-cables were closely related to a shorter distance from their perching sites to the shoreline. In other regions, the preference of these birds to perch in structures close to coastal and freshwater bodies has also been reported [40, 41, 88]. In the case of the CPCV, the 13 m-high lighting poles located between 15 and 60 m from the shoreline would offer the NECOs the best vantage points towards the sea. This, in turn, could favor the detection of feeding opportunities through the observation of other congeners or other predators that feed at sea [89, 90]. Because NECOs feed near their colonies (<2.5 km, [35]), their position on the pole-cables would make it possible to visualize much of the range of their feeding areas.

The high and positive linear correlation between DSZ:DS ratio and DSZ allows a clearer interpretation of the results. In this sense, closer distances from the pole-cables to the surf zone were associated with higher numbers of NECOs on these perching structures. This behavior could be because these areas are characterized by being quite biodiverse and promoting fish concentration [51, 91]. In fact, during the development of the study, some NECOs were observed swimming and diving among the waves of the surf zone, which agrees with previous observations in Costa Verde [42].

### Eco-friendly proposals for managing the environmental problem

Besides possible public health problems, traffic accidents or corrosion damage to vehicles and infrastructure, it has been evidenced throughout the study period that the feces of the NECOs do generate discomfort to people, esthetic problems (white spots of excrement on the road and sidewalk) and pestilence, as occurs in coastal cities of Chile [26]. These facts highlight the importance of proposing solutions to this environmental problem.

On the basis of the strategies to limit the interaction between cormorants and fishing activities [92], some of these could be adapted to the case of the NECOs in the CPCV. These strategies range from nonlethal deterrent methods to direct hunting. In some regions, the application of the latter to reduce populations of NECOs has been considered as a noneffective method [22, 93]. On the other hand, the use of nonlethal deterrents, such as sound stressors (e.g, sound-emitting bird scarers, noise-making devices such as cannons, guns and fireworks), has led to a habituation of the NECOs, and therefore to a null effectiveness [26, 93]. It should be mentioned that neither direct poaching nor noise-making devices (projectiles) would be adequate for the CPCV due to the possibility of harming people and because their application would be unethical. Visual stressors, such as silhouettes that emulate their predators, may also not be effective since no predators of this species are known in Costa Verde. The installation of perch deterrents (e.g., steel spikes, steel inverted-Y structures, dented triangular structures) on the lighting poles could reduce the perching activity of the NECOs, as has been reported with birds of prey in electrical transmission lines [94, 95] and with cormorants and gulls in oyster cages in Canada [96]. Nevertheless, experiences with NECOs on lighting poles on the coastal edge of the city of Antofagasta, Chile, have shown that this species rests without difficulty on these spikes and even bends them [26]. Taking into consideration both the preference of the NECOs to settle in areas near the shoreline and the surf zone, and the fact that in Costa Verde the nearest distances between the surf zone and the groynes are relatively short (range from 0 to 154 m), we propose the construction of structures with multiple perches located on the groynes, so as not to interfere with the activities of people on the beaches and let their feces fertilize the sea [97]. Simultaneously, decoys and vocalizations could be used to attract the NECOs to the new structures, which is effective for various seabird species [98, 99].

## Conclusions

This study is the first to elucidate the factors that influence the spatial distribution of a seabird species in coastal urban areas in Peru. The presence of NECOs on pole-cables is not unnoticed in Costa Verde since their droppings fall into the road, cars, sidewalks and passers-by, causing discomfort, pestilence and esthetic problems (white spots of excrement on the road and sidewalks). Despite these negative effects, the nuisance has not escalated to municipal action yet. However, with the development of new urban projects in Costa Verde (e.g., on February 27, 2020, an additional northern section of 2.7 km of the CPCV was inaugurated), the information of this research becomes increasingly relevant for better planning and reduction of conflicts between NECOs and people. The best management proposal that is offered based on the results of this study is to discourage these birds from using the poles and cables of the CPCV and at the same time relocate them to new perching structures on nearby groynes. It is recommended to continue with the monthly monitoring of the number of NECOs in the CPCV, studies of characterization of microbiota in their excrements, and pilot projects to evaluate the aforementioned management proposals. Likewise, it is important to update and increase the physical-environmental studies in Costa Verde in order to have a better understanding of this coastal ecosystem and its dynamics.

## Acknowledgments

This study is Sebastián Lozano-Sanllehi bachelor’s thesis for the degree of Environmental Engineer, Facultad de Ciencias Ambientales, Universidad Científica del Sur, Lima, Peru. We wish to thank Patricia Sanllehi, Mercedes Gaviria, Carolina Símbala, Keiji Aróstegui, Mariano Gutiérrez and Salvador Peraltilla for their help and suggestions for the data collection. A special thanks to Marcelo Stucchi for useful comments and discussions about Neotropic Cormorants in Costa Verde. We are also grateful to Bethany Clark for her support in the data analysis, and to Carolina Símbala and Diego Gonzáles for their comments and contributions in the design and making of some figures.

## Supporting Information Captions

**S1 Table. Number and type of lighting poles, length of the highway and number of groynes per district along the Circuito de Playas de la Costa Verde highway in Lima, Peru.**

**S1 File. Frequency distributions of the number of pole-cables used (red bars) or not used (blue bars) by Neotropic Cormorants (*Phalacrocorax brasilianus*) at 1000 h in the Circuito de Playas de la Costa Verde highway in Lima, Peru.**

**S2 Table. Statistical results of the fixed effects of the Poisson model: DS + DSZ:DS ratio + PG, better selected to explain the spatial distribution of the Neotropic Cormorants (*Phalacrocorax brasilianus*) in the Circuito de Playas de la Costa Verde highway in Lima, Peru.**

